# Regulation of SETD2 stability is important for the fidelity of H3K36me3 deposition

**DOI:** 10.1101/2020.05.21.100552

**Authors:** Saikat Bhattacharya, Jerry L. Workman

## Abstract

The histone H3K36me3 mark regulates transcription elongation, pre-mRNA splicing, DNA methylation, and DNA damage repair. However, knowledge of the regulation of the enzyme SETD2, which deposits this functionally important mark, is very limited. Here we show that the poorly characterized N-terminal region of SETD2 plays a determining role in regulating the stability of SETD2. This stretch of 1-1403 amino acids contributes to the robust degradation of SETD2 by the proteasome. Besides, the SETD2 protein is aggregate-prone and forms insoluble bodies in nuclei especially upon proteasome inhibition. Removal of the N-terminal segment results in the stabilization of SETD2 and leads to a marked increase in global H3K36me3 which, uncharacteristically, happens in a Pol II-independent manner. Thus, the regulation of SETD2 levels through proteasomal mediated decay is important to maintain the fidelity of H3K36me3 deposition.

## INTRODUCTION

The N-terminal tails of histones protrude from the nucleosome and are hotspots for the occurrence of a variety of post-translational modifications (PTMs) that play key roles in regulating epigenetic processes. H3K36me3 is one such important functionally characterized PTM. In yeast, this mark suppresses cryptic transcription from within the coding region of genes by preventing histone exchange [1]. In mammalian cells, it is involved in the recruitment of DNA repair machinery, in splicing and also, in establishing DNA methylation patterns by acting as a binding site for the enzyme DNMT3a [2][3][4][5][6]. Recent reports have emphasized the tumor-suppressive role of H3K36me3 in renal cancer especially, where the gene coding for the SETD2 histone methyltransferase is often deleted or mutated [7][8][9].

In yeast, the SET domain-containing protein Set2 (ySet2) is the sole H3K36 methyltransferase [10]. ySet2 interacts with the large subunit of the RNA polymerase II, Rpb1, through its SRI (**S**et2-**R**pb1 **I**nteraction) domain, and co-transcriptionally deposits H3K36me3 [11]. The deletion of the SRI domain from ySet2 abolishes both the Set2-RNA Pol II interaction and H3K36me3 methylation in yeast [12]. H3K36 methylation is a highly conserved histone mark and Set2 homologs are found in more complex eukaryotes [13]. These homologs share the conserved features like the AWS (**a**ssociated **w**ith **S**ET), SET [**S**u(var)3-9, **E**nhancer-of-zeste and **T**rithorax] and Post-SET domains that are required for the catalytic activity of the enzyme, and also, the protein-protein interaction domains such as the WW and the SRI. Notably, the mammalian homolog, SETD2, has a long N-terminal segment that is not present in ySet2. The function of this region has remained obscure [13][14].

Here we show that SETD2 is an inherently aggregate-prone protein and its N-terminal region regulates its half-life. This, in turn, is important for the fidelity of the deposition of the functionally important H3K36me3 mark. Our findings reveal that the previously uncharacterized N-terminal region of SETD2 is important in governing appropriate SETD2 function and activity.

## RESULTS

### SETD2 is robustly degraded by the ubiquitin-proteasome pathway

To understand the regulation of the SETD2 enzyme in human cells, we introduced a construct to express Halo- or GFP-tagged SETD2 FL (full length) under the control of CMV promoter in 293T cells. Strikingly the expression of GFP-SETD2 FL was barely detected post-72 hour of transfection although the expression of GFP in vector control transfected cells was robust [supplementary information S1a]. Also, RT-PCR revealed a marked increase in the transcript level of SETD2 suggesting a robust transcription from the constructs introduced [supplementary information S1b]. Similar results were obtained with Halo-SETD2 FL transfected cells in which the expression was not detected with the western blotting of whole-cell lysates with an anti-Halo antibody [Figure 1a].

**Figure 1:**
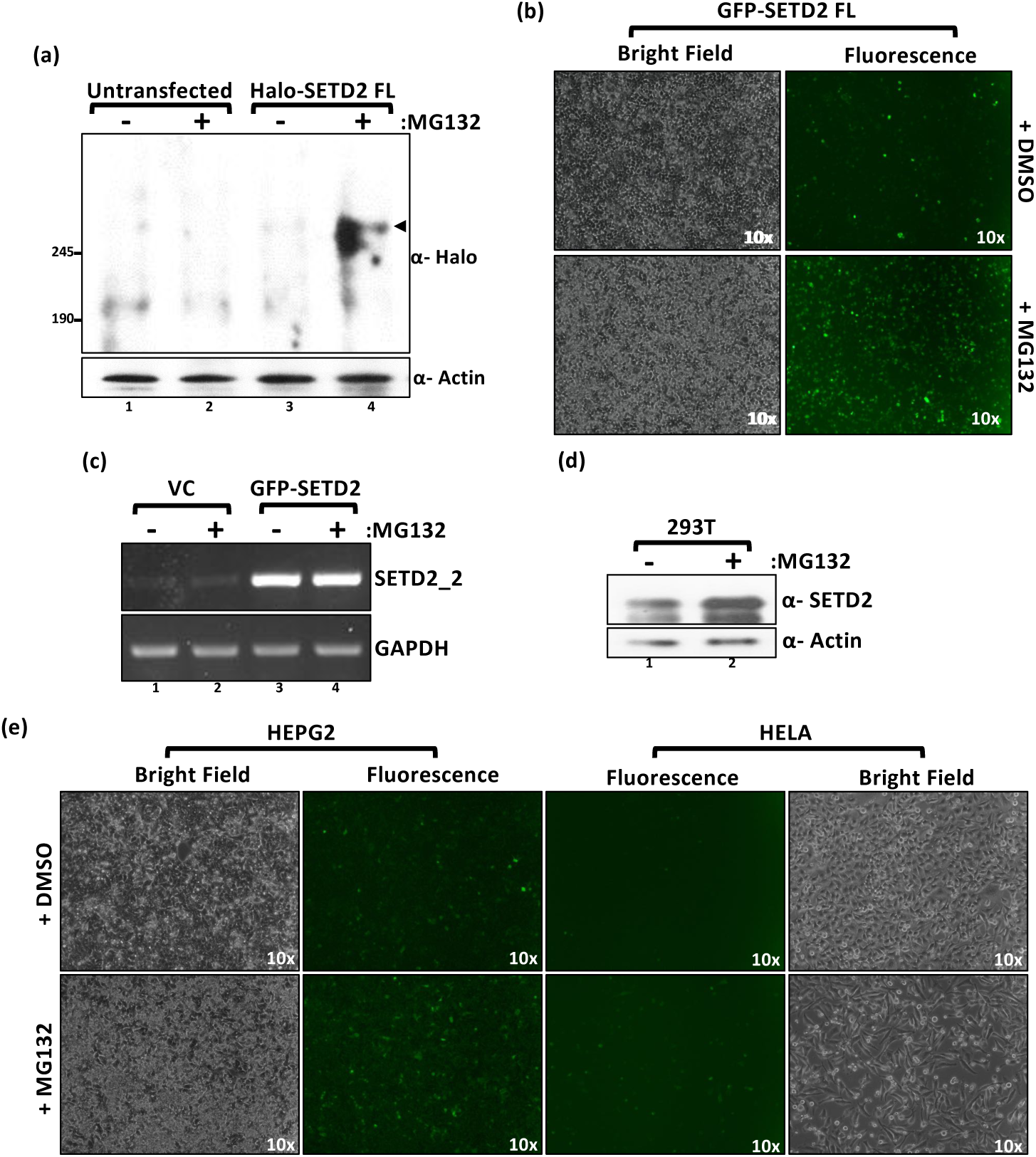
SETD2 is robustly degraded by the proteasome. (a) Western blot of whole-cell lysates probed with the depicted antibodies. Lysates of wild type 293T (untransfected) cells or expressing SETD2 full-length (FL) were prepared after 12 hrs of MG132 (10 µM) treatment. The expected band for the target protein is indicated by an arrow. (b) Microscopy images showing the effect of MG132 treatment on expression of GFP-SETD2 FL in 293T cells. (c) RNA was isolated from GFP-SETD2 FL transfected cells and RT-PCR was performed to check the transcript levels. GAPDH was used as a normalization control. VC-empty vector control. (d) Whole-cell lysates of 293T cells were prepared after 12 hrs of MG132 (10 µM) treatment and probed with the antibodies depicted. (e) Microscopy images showing the effect of MG132 treatment on the expression of GFP-SETD2 FL in the cell lines described.

Autophagy and the ubiquitin-proteasome system (UPS) are the major pathways for protein degradation in mammalian cells [15]. To investigate whether autophagy plays a role in SETD2 turn-over, 293T cells expressing GFP-SETD2 FL were treated with increasing concentration of the lysosome inhibitor chloroquine. Chloroquine treatment did not have an apparent effect on the GFP-SETD2 FL level [supplementary information S1c].

To test whether SETD2 is targeted for degradation by the UPS, SETD2 FL expression was checked after treating the cells with the proteasome inhibitor MG132. Proteasome inhibition led to an increase in the accumulation of SETD2. This was confirmed by western blotting of Halo-SETD2 FL expressing cells as well as microscopy of GFP-SETD2 FL [Figure 1a and b]. The addition of MG132 did not have a prominent effect on Halo or GFP expression [supplementary information S1d, e]. Also, RT-PCR did not reveal any change in the transcript abundance of both endogenous as well as exogenous SETD2 upon MG132 treatment, suggesting that the increased SETD2 protein abundance observed is due to protein stabilization [Figure 1c]. Importantly, an increase in the accumulation of the endogenous SETD2 was also observed by western blotting upon MG132 treatment of WT 293T cells, implying that degradation by the proteasome is not limited to the recombinant protein [Figure 1d]. Further, the expression of GFP-SETD2 FL was tested in HEPG2 and HELA cells. SETD2 behaved similarly in these cell lines. Very weak expression of GFP-SETD2 FL was observed which increased upon proteasome inhibition by MG132 treatment [Figure 1e].

Our results suggest that the short half-life of SETD2 is due to UPS mediated decay which is consistent with previous findings [16]. Also, this mechanism for SETD2 proteolysis is not cell line specific.

### The removal of N-terminus region stabilizes SETD2

Previously, the expression of yeast Set2 (ySet2) has been used in human cells to rescue H3K36me3 [17]. Hence, we next investigated the expression of Halo-ySet2 in 293T cells. Western blotting of whole-cell lysate with an anti-Halo antibody revealed robust expression of ySet2 [Figure 2a]. ySet2 is a well-characterized protein that is degraded by the proteasome in yeast [18]. The UPS and the architecture of the proteasome itself are conserved from yeast to mammals [19][20]. Consistent with that, our data shows that ySet2 responds to MG132 treatment in 293T cells [Figure 2a]. This suggests that ySet2 is targeted for degradation through UPS in human cells too although its expression level is much higher than that of SETD2 FL. SETD2 has an N-terminal region which is absent in ySet2 [Figure 2b]. We speculated that the disparity in expression between ySet2 and SETD2 FL could be due to the presence of this segment in SETD2.

**Figure 2:**
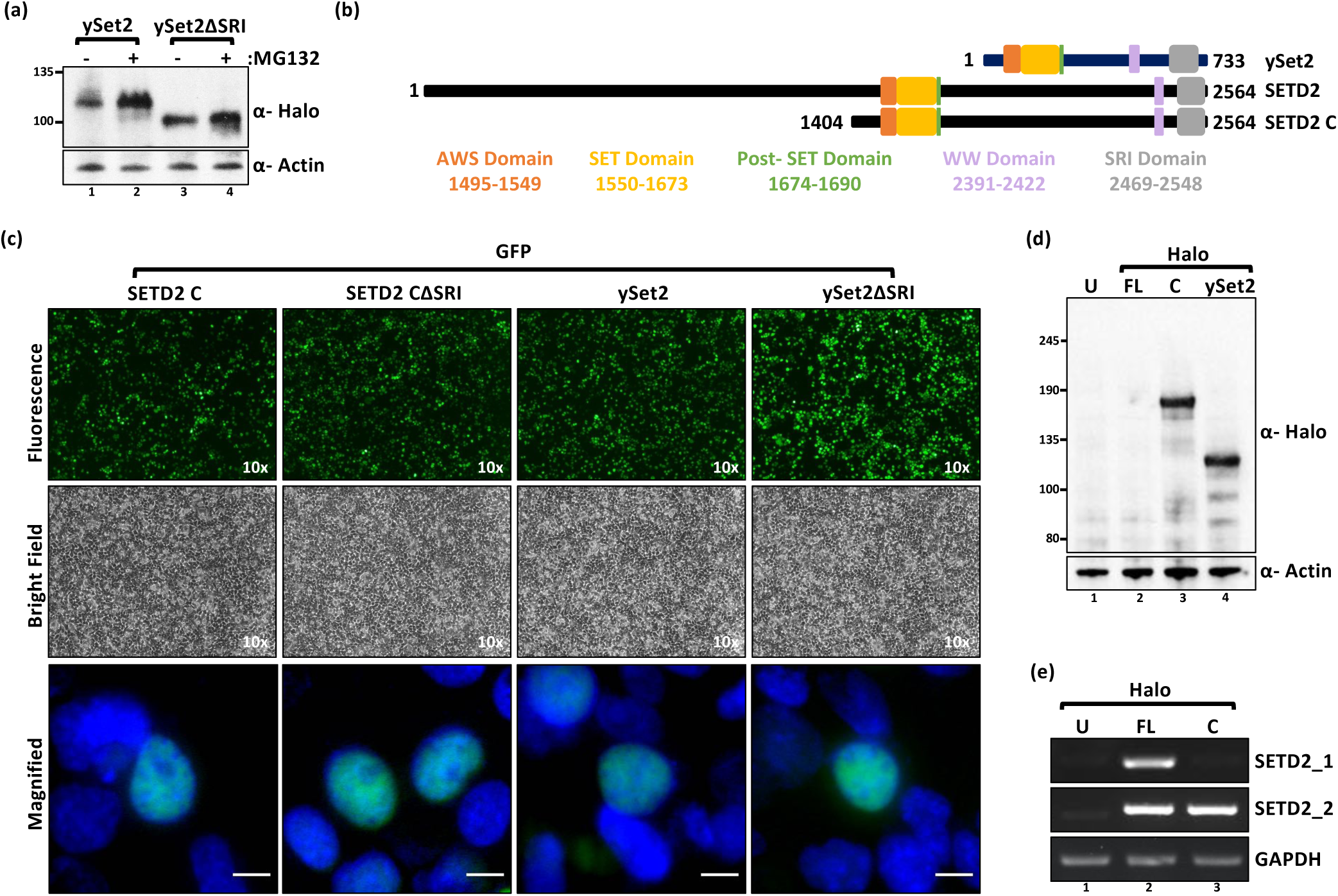
Removal of the N-terminal region of SETD2 leads to its stabilization. (a, d) Western blot of whole-cell lysates probed with the depicted antibodies. Lysates of 293T cells expressing Halo-ySet2 constructs were prepared after 12 hrs of MG132 (10 µM) treatment. U-untransfected. (b) Cartoon illustrating the known domains of yeast Set2 (ySet2), SETD2, and the SETD2 C-terminus region that shares the conserved domains with ySet2. (c) Microscopy images showing the expression and localization of GFP-SETD2 C and ySet2. The scale bar is 10 um. (e) RNA was isolated from transfected cells described in Figure 2d and RT-PCR was performed to check the SETD2 transcript levels. GAPDH was used as a normalization control. SETD2_1 primer pair binds to the N-terminal segment of SETD2 transcripts, whereas, SETD2_2 binds to the C-terminal region which is common in both SETD2 FL and C.

For this, we tested the expression level of a truncated mutant of SETD2, SETD2 C. Consistent with our hypothesis, microscopy of the GFP-tagged proteins revealed that the expression level of SETD2 C was comparable to that of ySet2 [Figure 2c]. Analysis of whole-cell lysate with anti-Halo western blotting revealed that the expression of SETD2 C was markedly higher than SETD2 FL and was comparable to ySet2 [Figure 2d]. RT-PCR using primers specific for SETD2 FL and SETD2 C transcripts confirmed that the transcripts were produced robustly [Figure 2e]. Also, the transcript levels of SETD2 FL and SETD2 C were comparable, suggesting that the differences observed in protein abundance are not due to the differences in the transcription of the two constructs but rather, might be due to the instability of the full-length SETD2 protein.

### Unlike the full-length protein, SETD2 fragments express robustly

We wanted to test whether a specific stretch of the N-terminal region of SETD2 causes its robust degradation. To test that, we made sub-fragments of the N-terminal region, 1-503 (Na) and 504-1403 (Nb), and tested their expression [Figure 3a]. Strikingly, western blotting of whole-cell lysates with an anti-Halo antibody revealed that both the fragments expressed robustly in 293T cells with their expression level similar to SETD2 C, whereas SETD2 FL could not be detected [Figure 3b]. Notably, microscopy analyses revealed that the fragment Nb was cytoplasmic, unlike Na and C which were nuclear [Figure 3c]. To test whether the localization of this fragment alters its stability, the C-myc nuclear localization signal (NLS) was added to GFP-Nb (Nb’) and the expression was checked. Western blotting of the whole-cell lysates revealed a reduced expression level of Nb’ as compared to Nb with continued sensitivity to MG132 treatment [Figure 3d]. Microscopy confirmed that the addition of NLS resulted in the nuclear translocation of Nb and also, reduced its expression level [Figure 3e]. Nevertheless, the expression of all the SETD2 fragments was very robust compared to SETD2 FL and continued to display sensitivity to MG132.

**Figure 3:**
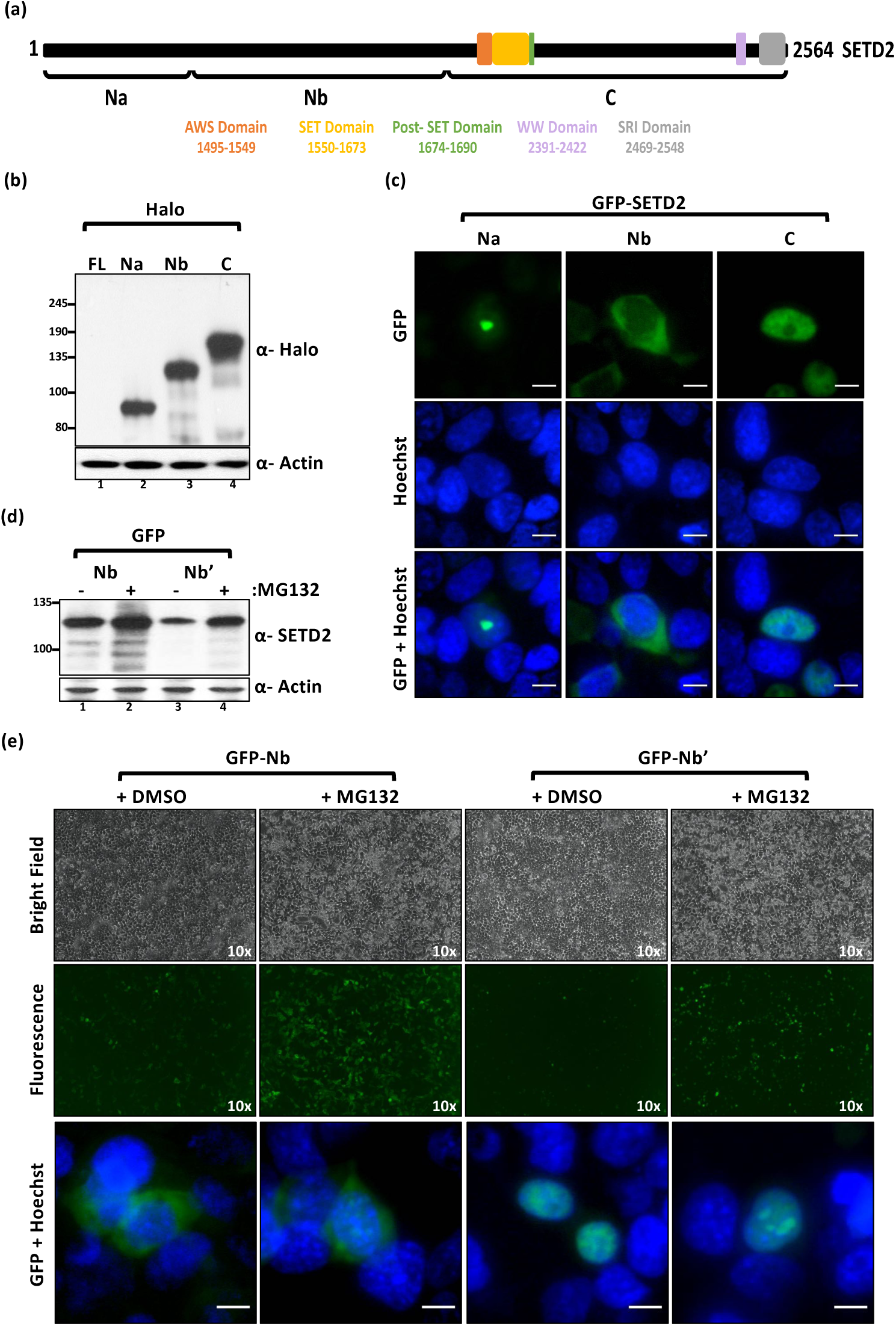
SETD2 fragments express robustly. (a) Cartoon illustrating the fragments of SETD2 along with its known domains. (b) Western blot of whole-cell lysates of 293T cells expressing Halo-SETD2 constructs probed with the depicted antibodies. (c, e) Microscopy images showing the localization of GFP-SETD2 fragments. The scale bar is 10 um. See the text for more details. (d) Western blot of whole-cell lysates of cells shown in figure 3e probed with the depicted antibodies.

From these experiments, no specific region emerged in the SETD2 protein that is particularly targeted for UPS-mediated decay.

### SETD2 forms nuclear puncta

Microscopy revealed that GFP-SETD2 Na is nuclear and formed puncta [Figure 3c]. This was surprising because this segment lacks an NLS with a significant score as per NLS mapper prediction (http://nls-mapper.iab.keio.ac.jp/cgi-bin/NLS_Mapper_form.cgi) [21]. We decided to characterize the NLS of SETD2 to better understand the unexpected localization of the SETD2 fragments. To this end, the localization of a series of GFP-SETD2 fragments was checked using fluorescence microscopy (data not shown). Of all the SETD2 fragments tested, the data revealed the presence of putative NLS in three fragments of SETD2: 967-1690, 1964-2263, and 2423-2564 [Figure 4a]. This was consistent with the NLS mapper prediction that revealed the presence of NLS in each of these segments [supplementary information S2]. To validate these NLS, site-directed mutagenesis was performed to mutate the K and R residues to A and the disruption of the nuclear localization of the mutated GFP-SETD2 fragments was confirmed by microscopy [Figure 4a, supplementary information 2b]. Further, to validate that the full-length SETD2 has only these three NLS, site-directed mutagenesis was performed to mutate these NLS one by one [Figure 4b]. The cytoplasmic localization of the GFP-SETD2 FL mutant, in which all the three NLS were disrupted, confirmed that SETD2 has three NLS [Figure 4b]. Importantly, this also shows that the SETD2 fragment Na forms nuclear puncta without an NLS.

**Figure 4:**
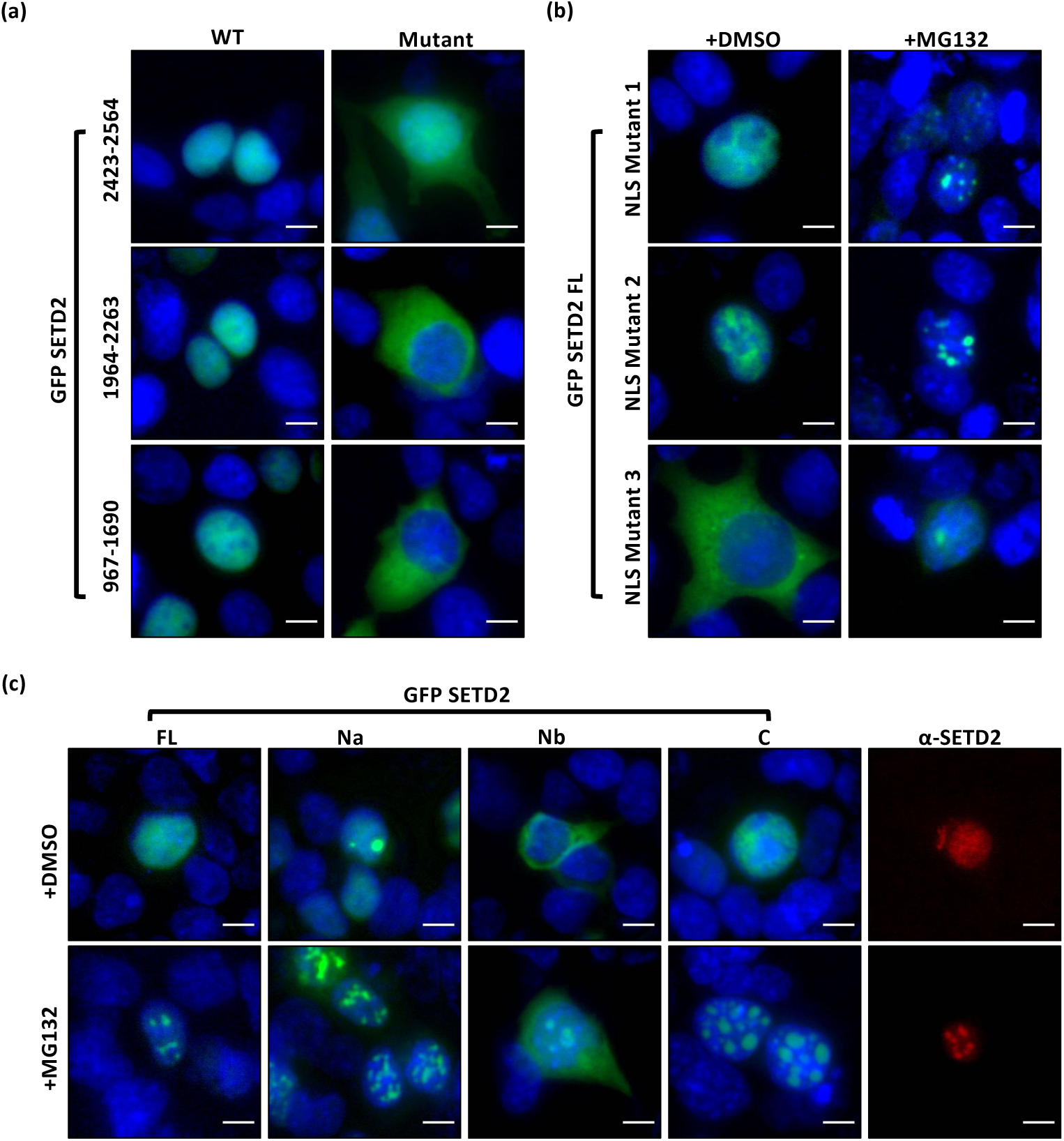
SETD2 forms nuclear puncta. (a, b) Microscopy images showing localization of GFP-SETD2 fragments that have putative NLS and the NLS mutants. See the text for more details. (c) Microscopy images showing the localization of SETD2 and its fragments. The scale bar is 10 um.

To test whether the full-length SETD2 protein also behaves similarly, the localization of NLS mutants of SETD2 was tested on MG132 treatment. Strikingly, all the mutants exhibited the formation of nuclear puncta [Figure 4b]. To characterize the aggregate-prone tendency of SETD2 further, 293T cells expressing GFP-SETD2 truncations (described in Figure 2) were observed under the microscope with or without MG132 treatment. Remarkably, all the SETD2 fragments responded to the treatment with MG132 and demonstrated a marked increase in the formation of puncta [Figure 4c]. Notably, the completely cytoplasmic fragment Nb showed a pan-cellular distribution with a tendency to form puncta upon MG132 treatment similar to SETD2 FL NLS Mutant 3. Furthermore, puncta were also formed by WT GFP-SETD2 FL protein suggesting that it might not be an artifact caused due to protein misfolding resulting from the truncations or mutations [Figure 4c]. To test whether the endogenous SETD2 behaves similarly, immunofluorescence of 293T cells was performed with an anti-SETD2 antibody. Importantly, proteasomal inhibition caused speckle-like staining, revealing that endogenous SETD2 also forms puncta [Figure 4c]. This observation, together with the fact that despite very weak expression, SETD2 FL is aggregate-prone, suggests that aggregation is an intrinsic property of the SETD2 protein that is exacerbated by an increased protein abundance.

### SETD2 forms ubiquitinated insoluble aggregates

Exogenously expressed as well as the endogenous SETD2 formed puncta, especially upon MG132 treatment, that are reminiscent of inclusion bodies formed by aggregated proteins. The most striking pattern was observed with the fragment SETD2 Na that formed distinct puncta even in the absence of proteasome inhibition. As aggregates are often ubiquitinated, to confirm that the puncta formed by SETD2 are aggregated structures, the colocalization of RFP-ubiquitin with GFP-SETD2 Na was tested. Clear colocalization was observed suggesting that SETD2 Na puncta are indeed ubiquitinated aggregate structures [Figure 5a]. GFP-SETD2 Na did not colocalize with RFP-Fibrillarin indicating that the puncta are not nucleolar [Figure 5a]. To biochemically substantiate that SETD2 is ubiquitinated, Halo-C was affinity-purified from HEK293T cells co-expressing HA-ubiquitin with or without MG132 treatment. The purified proteins were then resolved on a gel and analyzed for the presence of ubiquitination. Western blotting with an anti-HA antibody revealed that SETD2 C was indeed ubiquitinated [Figure 5b].

**Figure 5:**
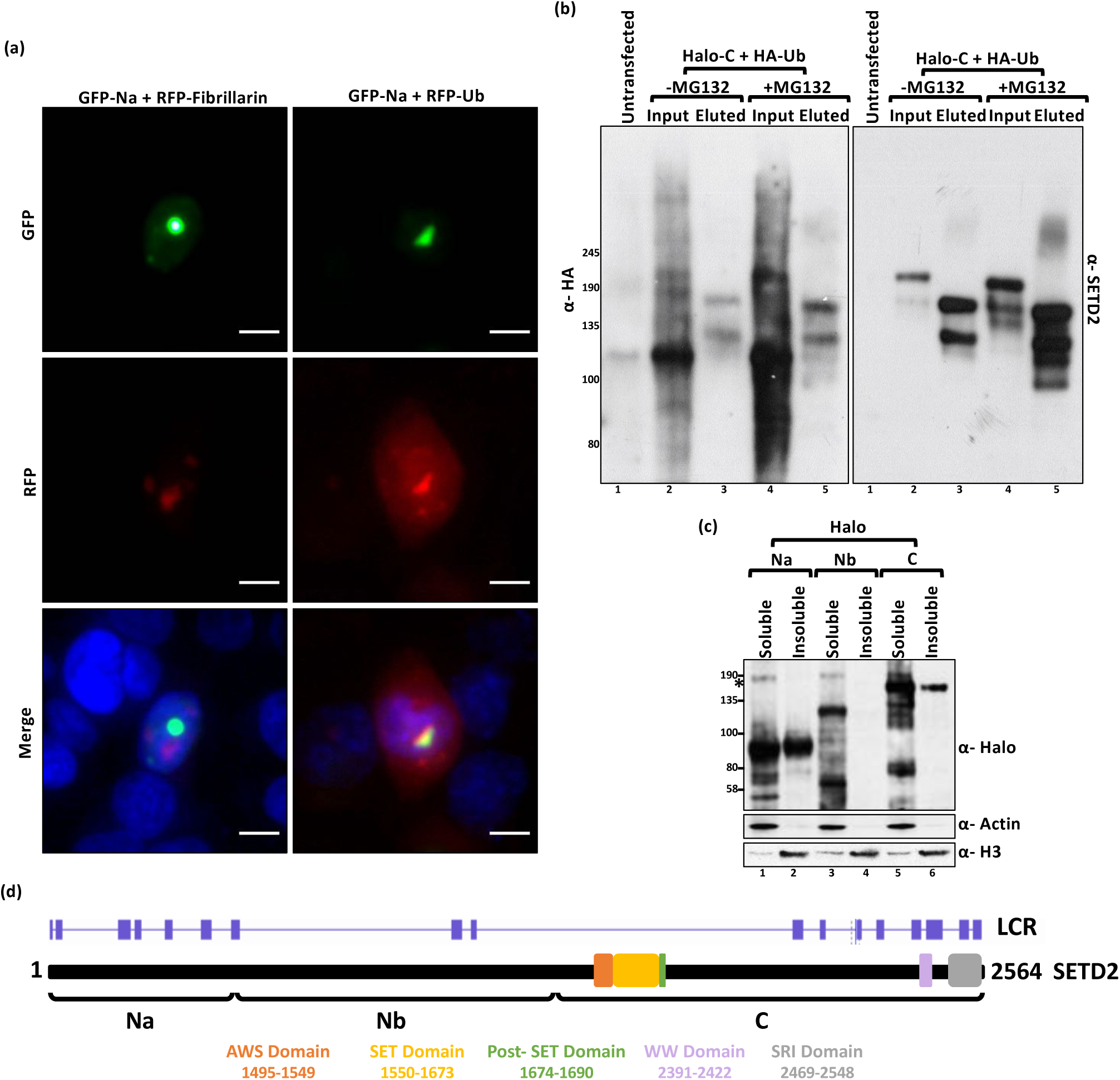
SETD2 forms insoluble ubiquitinated aggregates. (a) Microscopy images showing the colocalization of GFP-SETD2 Na fragment with RFP-Fibrillarin or RFP-Ub. The scale bar is 10 um. (b) Halo purification was performed from extracts of 293T cells co-expressing Halo-C and HA-Ub. Eluted and 0.5% of Input samples were resolved on a gel and probed with the depicted antibodies. (c) Soluble and insoluble fractions were separated from 293T cells expressing Halo-SETD2 fragments, resolved on a gel, and probed with the depicted antibodies. *-non-specific. (d) Cartoon depicting the predicted low-complexity regions (LCRs) in SETD2. The prediction was performed using the webserver at http://repeat.biol.ucy.ac.cy/fgb2/gbrowse/swissprot/.

To confirm further that SETD2 forms aggregates, we checked the solubility of Halo-SETD2 fragments Na, Nb, and C. 293T cells expressing the Halo-SETD2 fragments were lysed, their soluble and insoluble fractions separated and afterward, analyzed by western blotting with an anti-Halo antibody. Correlating with the microscopy observations, SETD2 Na that formed spontaneous puncta was highly insoluble; C was insoluble to a lesser extent [Figure 5c]. The segment Nb was soluble [Figure 5c]. Interestingly, the data shows that the different regions of the same protein have different aggregation propensity under similar experimental conditions and are not merely a consequence of expression from a strong CMV promoter. Furthermore, the presence of low-complexity regions (LCR) in a protein is linked to its aggregation propensity [22][23][24]. Hence, we analyzed the SETD2 protein sequence for the presence of LCR. Strikingly, the solubility profile observed for the SETD2 fragments correlated very well with the LCR distribution of the protein [Figure 5d].

Collectively, our data show that SETD2 forms aggregated ubiquitinated puncta (discussed in the Discussion section).

### Removal of the N-terminal region of SETD2 leads to a marked increase in global H3K36me3 levels

We found that the removal of the N-terminal region of SETD2 leads to the stabilization of the remaining portion that shares conserved domains with ySet2. Next, we investigated whether the removal of the N-terminal region affects the catalytic activity of SETD2. To check the activity of the exogenously introduced SETD2, *setd2Δ* (KO) 293T cells were used. Consistent with the role of SETD2 as the sole H3K36me3 depositor in humans, in the KO cells the H3K36me3 mark was not detected in the whole-cell lysates by immunoblotting [Figure 6a]. Next, constructs to express Halo-tagged SETD2 FL or SETD2 C were introduced in the KO cells by transfection. 72 hours post-transfection, whole-cell lysates were prepared and analyzed by western blotting. The expression of the empty vector (VC) did not rescue H3K36me3 as expected [Figure 6a]. Strikingly, the expression of SETD2 C in KO cells led to a marked increase in the H3K36me3 level as compared to the rescue with SETD2 FL [Figure 6a]. The other two H3K36 methyl marks, H3K36me1 and H3K36me2, largely remained unchanged [Figure 6a].

**Figure 6:**
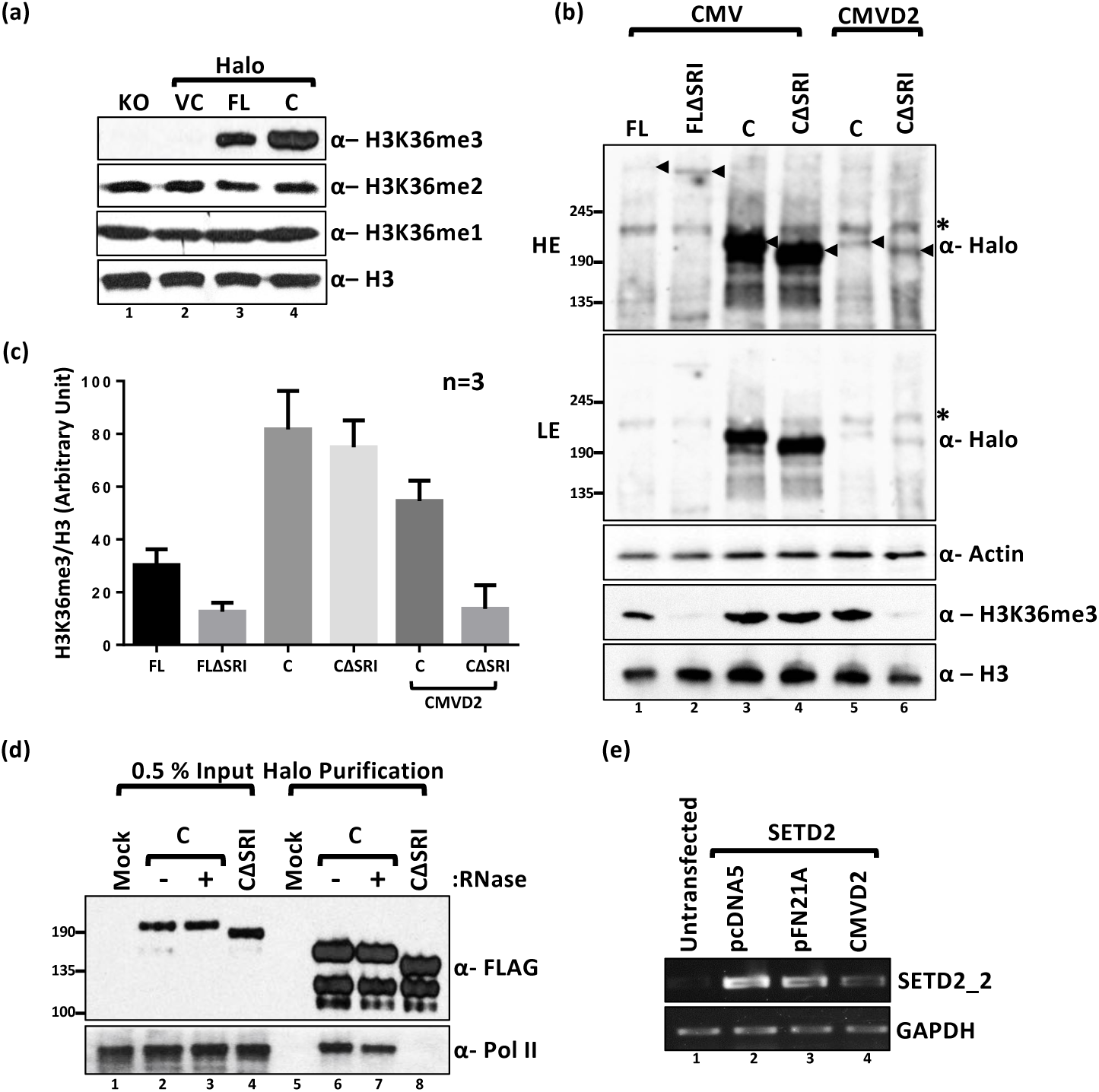
SETD2 has reduced Pol II dependency at high expression levels. (a) Western blot of whole-cell lysates probed with the depicted antibodies. setd2Δ 293T (KO) cells were transfected with Halo-vector control (VC), SETD2 full-length (FL) or SETD2 C (C), and the lysates were prepared 72 hrs post-transfection. (b) Western blot of whole-cell lysates of 293T cells expressing Halo-SETD2 constructs, under the control of either CMV or CMVD2, probed with the depicted antibodies. The expected bands are marked by arrows. *-non-specific. (c) Bar graph showing H3 normalized H3K36me3 signal intensity of data depicted in Figure 6b. The data plotted are from three independent biological replicates. (d) Halo purification was performed from extracts of 293T cells expressing Halo-SETD2 C. Input and eluted samples were resolved on a gel and probed with the depicted antibodies. (e) RNA was isolated from 293T cells expressing SETD2 from different vectors and RT-PCR was performed to check transcript levels. GAPDH was used as a normalization control.

Thus, the removal of the N-terminal segment leads to a marked increase in the global H3K36me3 level. Also, these experiments demonstrate that in the absence of the N-terminal region, SETD2 retains its histone methyltransferase activity.

### At high cellular levels, SETD2 has a reduced RNA pol II dependency for H3K36me3 deposition

Studies conducted in yeast have revealed that the deposition of the H3K36me3 mark is strictly dependent on the ySet2-Pol II association [12]. We were curious whether the marked increase in the global H3K36me3 level upon SETD2 C expression happens in an RNA Pol II-dependent manner.

To test this, Halo-SETD2 constructs without the SRI domain were introduced in *setd2Δ* 293T cells. Similar to the findings for ySet2 in yeast, removal of the SRI domain from full-length SETD2 protein (FLΔSRI) led to a marked decrease in H3K36me3 levels as compared to the FL [Figure 6b, c]. Strikingly though, removal of the SRI domain from SETD2 C (CΔSRI) had a very marginal effect on the H3K36me3 levels [Figure 6b, c]. To confirm that the removal of the SRI domain leads to the abolishment of SETD2-Pol II interaction, Halo-FLAG-SETD2 C and Halo-FLAG-SETD2 CΔSRI were affinity purified from 293T extracts using Halo ligand-conjugated magnetic resin. Elution of proteins purified using this technique involves cleaving the Halo tag with TEV protease, leaving the FLAG epitope which can be detected on the eluted bait by immunoblotting [Figure 6d]. Immunoblotting with an anti-Pol II antibody confirmed that the deletion of the SRI domain from SETD2 leads to the abolishment of SETD2-Pol II interaction [Figure 6d].

We wondered whether the decreased dependency on Pol II interaction for the H3K36me3 activity of SETD2 C is due to the loss of a possible autoinhibition by the N-terminal region of SETD2 or is due to the increased expression of SETD2 C fragment as compared to the full-length protein. To address these possibilities, Halo-SETD2 C constructs under the control of the CMVD2 promoter were introduced in *setd2Δ* cells. CMVD2 promoter is a truncated form of CMV and exhibits a much weaker activity. The weaker activity of the CMVD2 promoter was confirmed by RT-PCR [Figure 6e]. The reduced expression of SETD2 C under the regulation of CMVD2 promoter was verified by analyzing whole-cell lysate with an anti-Halo antibody [Figure 6b]. Notably, analysis of H3K36me3 revealed that the RNA Pol II dependency of SETD2 C was restored at a reduced expression level as SETD2 CΔSRI did not exhibit much activity when expressed using the CMVD2 promoter [Figure 6b, c].

We conclude that at high cellular levels, SETD2 has a reduced RNA pol II dependency for H3K36me3 deposition.

## DISCUSSION

In recent years many reports have highlighted the adverse effect of the loss of H3K36me3 that occurs upon SETD2 deletion. Here we show that the other end of the spectrum can also be detrimental as an excess of SETD2 can lead to inadvertent consequences. Previously, SETD2 has been reported to be regulated by the proteasome [16]. We reveal that the absence of SETD2 proteolysis results in a Pol II independent H3K36me3 deposition and protein aggregation. Our work illustrates the importance of the N-terminal segment of SETD2, which has been a mystery, in maintaining the requisite intracellular amount of the protein.

### Regulation of SETD2 half-life is important for regulating its function

Altered protein half-life can lead to abnormal development and diseases such as cancer and neurodegeneration [25]. Considering the role of H3K36me3 in a variety of important cellular processes, it is reasonable that regulating the activity of the methyltransferase responsible for the deposition of this mark is important. The previously uncharacterized N-terminal region, which is absent in ySet2, plays an important role in SETD2 regulation. The gain or loss of protein segments may be an important contributor to the degradation rate of proteins during evolution [26]. The differences in half-lives between homologs might be needed to adjust for the differences in the mechanism of deposition of H3K36 methylation. For instance, unlike in yeast where ySet2 performs all three states of methylation of H3K36, SETD2 does not appear to be majorly responsible for me1 and me2 deposition. Therefore, even with a shorter half-life, SETD2 might be able to do the required H3K36me3 deposition. Some evidence for this is provided by studies on human cancers that show that total H3K36me3 levels are not significantly impacted by a monoallelic loss of SETD2 [27][28].

### SETD2 over-abundance might have inadvertent consequences

We found that SETD2 is an inherently aggregate-prone protein and its aggregation tendency positively correlated with the presence of LCRs. Aggregation induced by LCRs can be detrimental and there are more than 20 genetic disorders linked to the expansion of trinucleotide repeats within coding sequences that generates LCRs. The uncontrolled expansion of CAG triplets that leads to polyQ tracts is associated with Huntington disease and several ataxias [29]. Overaccumulation of SETD2 can lead to the formation of stable and insoluble protein aggregates impeding the normal function of other proteins. SETD2 aggregation might lead to the co-aggregation of other proteins leading to their inactivation and proteotoxic stress. Ubiquitinated aggregates like the ones formed by SETD2 can directly inhibit or clog proteasomes [30][31]. As aggregation is a concentration-dependent phenomenon, possibly the robust degradation mechanism in cells ensures that SETD2 is kept at levels that maintain its solubility and activity. Interestingly, SETD2, aka, HYPB was initially identified in a screen to find interactors of the aggregate prone protein Huntingtin (Htt) [32]. It is possible that the reported Huntingtin-SETD2 interaction was due to the aggregation propensities of these proteins. Interestingly, the N-terminus of SETD2 (SETD2 Na) behaves very similarly to the polyQ containing N-terminus of Huntingtin protein. Like mutant Htt, SETD2 Na localizes to the nucleus in the absence of an NLS and forms spontaneous puncta characteristic of aggregated proteins [33]. The nucleoplasm promotes the formation of such aberrant and insoluble protein aggregates due to the strong crowding forces from highly concentrated macromolecules (approximately 100 mg/ml) [34]. It will be interesting to investigate in the future whether the aggregate prone tendency of SETD2 enables it to form a part of RNA-granules and regulate transcription.

### Pol II association is required for enhancing SETD2 activity

Our data shows that when the expression level of SETD2 is high, it has reduced dependency on RNA Pol II association for H3K36me3 deposition. Possibly the interaction with Pol II is required for the activation of SETD2 enzymatic activity. When present in its normal cellular amounts, SETD2 is not active enough to lead to robust H3K36me3 deposition without the Pol II association. Pol II association enhances SETD2 activity and hence, although SETD2 FL protein is barely detectable, the rescue of H3K36me3 can be readily seen in the *setd2Δ* cells. At high abundance, despite its low activity, a robust H3K36me3 deposition is seen likely due to the increased copy number of SETD2 protein in cells. Recent studies in yeast have also challenged the notion that the Pol II association is required for chromatin recruitment of Set2 [35]. The study found that the engagement of Set2 and Pol II through the SRI domain is rather required for the activation of Set2. Pol II independent deposition of H3K36me3 can result in the mistargeting of important epigenetic regulators. The PWWP domain-containing proteins DNMT3a, MutSα, and MORF depend on H3K36me3 for proper recruitment [4][5][36][37]. It is important to note that besides histone H3, SETD2 also has non-histone targets like tubulin, the methylation of which is important for metaphase transition [38]. Hence, the consequences of an increase in SETD2 abundance might not be limited to changes in histone methylation.

## MATERIALS AND METHODS

### Plasmids

SETD2-HaloTag® human ORF in pFN21A was procured from Promega. Deletion mutants of SETD2 were constructed by PCR (Phusion polymerase, NEB) using full-length SETD2 as a template and individual fragments were cloned. All constructs generated were confirmed by sequencing. SETD2-GFP, mRFP-Ub, pLenti puro HA-Ubiquitin, pTagRFP-C1-Fibrillarin and pCDNA3-ySet2 were procured from Addgene.

### Cell line maintenance and drug treatment

The cell lines used in this study (HEK293T, HEPG2 and, HELA) were procured from ATCC. Cells were maintained in DMEM supplemented with 10% FBS and 2 mM L-glutamine at 37 °C with 5% CO_2_. MG132 (Sigma) was added at a final concentration of 10 μM for 12 hours. Chloroquine (Sigma) treatment was done as indicated in the text. Transfections were performed at cell confluency of 40% using Fugene HD (Promega) using a ratio of 1:4 of the plasmid (µg) to transfection reagent (µl).

### Isolation of total RNA and PCR

Total RNA was extracted from cells as per the manufacturer’s (Qiagen) instructions. It was further treated with DNaseI (NEB) for 30 min at 72 °C to degrade any possible DNA contamination. RNA (2 μg) was subjected to reverse transcription using the QScript cDNA synthesis mix according to the manufacturer’s instructions. cDNAs were then amplified with the corresponding gene-specific primer sets. For RTPCR, PCR was conducted for 24 cycles using the condition of 30 s at 94 °C, 30 s at 60 °C and 30 s at 72 °C. The PCR products were analyzed on a 1% agarose gels containing 0.5 μg/ml ethidium bromide. The sequence of oligos is in supplementary information S3.

### Histone isolation and immunoblot analysis

Histones were isolated and analyzed as described previously [39]. For immunoblotting, histones were resolved on 15% SDS–polyacrylamide gel, transferred to PVDF membrane and probed with antibodies. Signals were detected by using the ECL plus detection kit (ThermoFisher).

### Antibodies

H3K36me3 (CST, 4909S), H3K36me2 (Active Motif, 39255), H3K36me1 (Abcam, ab9048), H3 (CST, 9715S), Halo (Promega, G9211), β-actin (Abcam, ab8224), SETD2 (Abclonal, A3194), FLAG (Sigma-Aldrich, A8592), Pol II (Abcam, ab5095).

### Cell Fractionation

To prepare soluble and insoluble extracts, 293T cells were washed with 1xPBS, collected by centrifugation, and resuspended in lysis buffer (50 mM Tris, pH 7.5, 350 mM NaCl, 1% Triton-X 100, 0.1% Na-deoxycholate and a protease inhibitor mix). The lysed cells were centrifuged at 13,000 rpm for 20 min. The supernatant was collected as the soluble fraction. The pellet was washed with lysis buffer containing 600 mM NaCl). The remaining insoluble pellet following another centrifugation was resuspended in Laemmli buffer (Biorad) and solubilized by sonication on ice.

### Affinity purification

293T cells expressing the protein of interest were harvested in 1xPBS and collected by centrifugation. The cells were lysed by resuspending in lysis buffer (50 mM Tris, pH 7.5, 150 mM NaCl, 1% Triton-X 100, 0.1% Na-deoxycholate and a protease inhibitor cocktail). The lysed cells were centrifuged at 13,000 rpm for 20 min. The supernatant was collected and diluted 1:3 by adding dilution buffer (1x PBS, pH 7.5 with 1mM DTT and 0.005% NP-40). The diluted lysate was added to 100 µl of pre-equilibrated Magne® HaloTag® Beads (Promega, G7282) and incubated overnight on a rotator at 4 °C. The beads were then washed with wash buffer (50 mM Tris-HCL, pH 7.5, 300 mM NaCl, 0.005% NP40 and 1mM DTT. 2 µl AcTEV (ThermoFisher, 12575015) protease was used per 100 µl of elution buffer.

### Immunofluorescence

293T cells were plated onto glass coverslips in a 6-well plate. Cells were washed with 1x PBS and fixed in 4% paraformaldehyde for 20 min at 37 °C. Cells were then washed three times with cold 1x PBS and permeabilized for 5 mins with 1x PBS containing 0.2% Triton X-100. Permeabilized cells were then blocked for 30 min with blocking buffer (3% BSA and 0.1% Triton-X in 1x PBS). Cells were stained with primary antibodies against SETD2 (1:1,000; Abclonal) for 1 hr at room temperature. A secondary antibody conjugated with AlexaFluor 568 was applied for 1 hr at room temperature. All the images depicted in the same panel were captured using the same settings.

## ACKNOWLEDGMENT

The authors are grateful to Dr. Kimryn Rathmell, Vanderbilt Institute for Infection, Immunology and Inflammation from providing SETD2 knock out 293T cells. The authors would like to thank the members of the Workman lab for their critical suggestions to improve the manuscript.

## AUTHOR CONTRIBUTION

S.B. conceptualized the work, designed and performed the experiments. S.B. wrote the manuscript. J.L.W conceived the idea of the work, provided supervision, acquired funding and revised the manuscript.

