## supplementary information for "Regulation of SETD2 stability is important for the fidelity of H3K36me3 deposition"

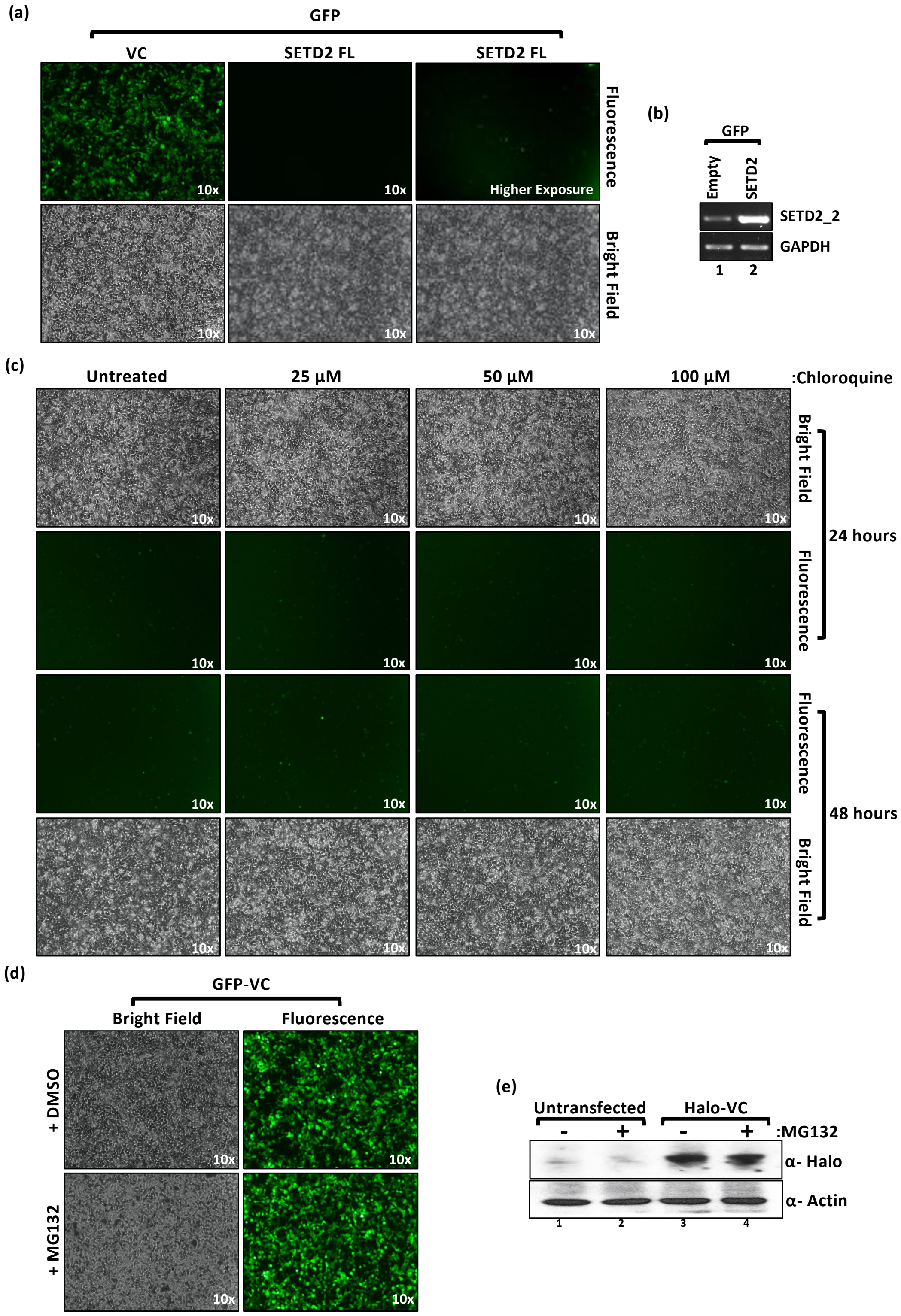

(a)
GFP
VC
SETD2 FL
SETD2 FL
10x
10x
Higher Exposure
Fluorescence
Bright Field
(b)
GFP
Empty
SETD2
SETD2_2
GAPDH
1
2
(c)
Untreated
25 µM
50 µM
100 µM
:Chloroquine
Bright Field
Fluorescence
24 hours
Fluorescence
Bright Field
48 hours
(d)
GFP-VC
Bright Field
Fluorescence
+ DMSO
+ MG132
(e)
Untransfected
Halo-VC
-
+
-
+
:MG132
α- Halo
α- Actin
1
2
3
4

**Supplementary information S1:** (a) Microscopy images showing expression of GFP constructs in 293T cells. The 'Higher Exposure' image is included to more clearly depict expression of the protein. (b) RNA was isolated from transfected cells described in (a) and RT-PCR was performed to check transcript levels. GAPDH was used as a normalization control. (c) Microscopy images showing the effect of chloroquine treatment on expression of GFP-SETD2 FL in 293T cells. 24 hours post-transfection, chloroquine was added to the culture media at the concentrations and for the time periods depicted. (d) Microscopy images showing effect of MG132 treatment on expression of GFP in 293T cells. (e) Western blot of whole-cell lysates probed with the depicted antibodies. Lysates of wild type 293T (untransfected) cells expressing Halo-vector control (VC) were prepared after 12 hrs of MG132 (10 µM) treatment.

(a)

| Predicted monopartite NLS | | |
|---|---|---|
| Position | Sequence | Score |
| 1400 | DRGPLKKRRQEIE | 8.5 |
| 1401 | RGPLKKRRQEI | 8.5 |
| 1403 | PLKKRRQEIE | 8 |

| Predicted bipartite NLS | | |
|---|---|---|
| Position | Sequence | Score |
| 1400 | DRGPLKKRRQEIESDSESDGELQDRKKVRVE | 6.5 |
| 1400 | DRGPLKKRRQEIESDSESDGELQDRKKVRVE | 5.2 |
| 1400 | DRGPLKKRRQEIESDSESDGELQDRKKVRVEVE | 5.3 |
| 1401 | RGPLKKRRQEIESDSESDGELQDRKKVRVE | 10.1 |
| 1401 | RGPLKKRRQEIESDSESDGELQDRKKVRVE | 6.5 |
| 1401 | RGPLKKRRQEIESDSESDGELQDRKKVRVEVE | 5.7 |
| 1401 | RGPLKKRRQEIESDSESDGELQDRKKVRVE | 5.6 |
| 2071 | NKEKRKRRSSLSPPSSAYERGTKRPDD | 6 |
| 2072 | KEKRKRRSSLSPPSSAYERGTKRPDD | 7.7 |
| 2089 | ERGTKRPDDRYDTPTSKKKVRIK | 5.6 |
| 2450 | KASKKPKTAEADTSSELAKKSKEVFRKEMS | 4.3 |

(b)

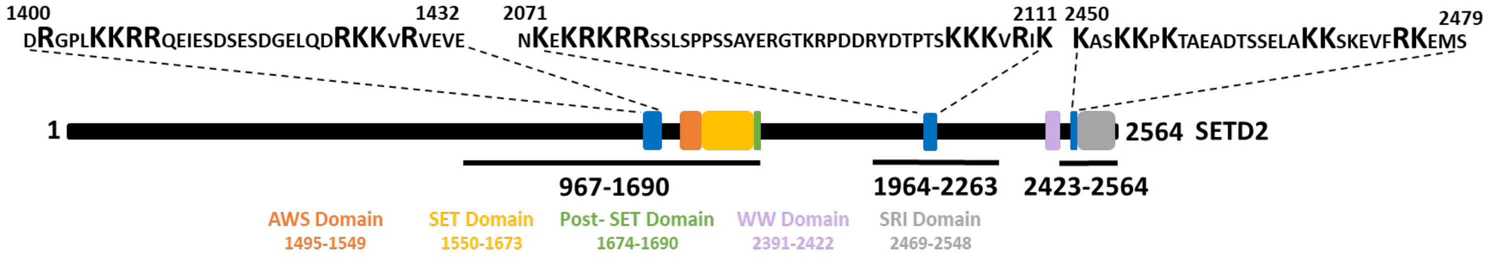


**Supplementary information S2:** (a) Position, sequence and score of putative NLS in SETD2 based on NLS Mapper prediction. (b) Cartoon illustrating the position and sequence of putative NLS based on NLS Mapper prediction. The highlighted K (lysine) and R (arginine) were mutated to A (alanine) to disrupt NLS in SETD2 fragments 967-1690, 1964-2263 and 2423-2564 as well in the full-length SETD2.

| Oligo | Sequence (5’-3’) |
|---|---|
| SETD2_1_F | GGGAAGGATGAAGAAATTCCAG |
| SETD2_1_R | CCCTAGATCCTCACTTTTTAAAGG |
| SETD2_2_F | AAGAAGCTCCCTCTCACCAC |
| SETD2_2_R | GATCCACATAGGCCTGCATG |
| GAPDH _F | TTCGACAGTCAGCCGCATCTTCTT |
| GAPDH _R | CAGGCGCCCAATACGACCAAATC |

**Supplementary information S3:** Sequence of oligos used to perform RT-PCR and knockdown.
